# Outer layer of Vb neurons in medial entorhinal cortex project to hippocampal dentate gyrus in mice

**DOI:** 10.1101/2023.11.06.565866

**Authors:** Naoki Yamamoto, Jun Yokose, Kritika Ramesh, Takashi Kitamura, Sachie K. Ogawa

## Abstract

Entorhinal cortical (EC)-hippocampal (HPC) circuits are crucial for learning and memory. Although it was traditionally believed that superficial layers (II/III) of the EC mainly project to the HPC and deep layers (V/VI) receive input from the HPC, recent studies have highlighted the significant projections from layers Va and VI of the EC into the HPC. However, it still remains unknown whether Vb neurons in the EC provide projections to the hippocampus. In this study, using a molecular marker for Vb and retrograde tracers, we identified that the outer layer of Vb in the medial EC (MEC) neurons directly project to both dorsal and ventral hippocampal DG, with a significant preference for the ventral DG. In contrast to the distribution of DG-projecting Vb cells, anterior thalamus-projecting Vb cells are distributed through the outer to the inner layer of Vb. Furthermore, dual tracer injections revealed that DG-projecting Vb cells and anterior thalamus-projecting Vb cells are distinct populations. These results suggest that the roles of MEC Vb neurons are not merely limited to the formation of EC-HPC loop circuits, but rather contribute to multiple neural processes for learning and memory.

## Main

In humans and rodents, the entorhinal cortical (EC)-hippocampal (HPC) networks are crucial for the formation and recall of episodic memory (1). Identification of cell-type specific projections in the EC-HPC networks is critical for understanding the neural process and the computations underlying learning and memory. The hippocampus can be divided into the dentate gyrus (DG), CA3, CA2, and CA1 regions (2, 3). It has been considered that the superficial layers (II/III) of EC mainly project to the HPC, while the deep layers (V/VI) receive the input from the HPC to provide telencephalic projections (4). For example, Reelin^+^ cells in layer II of the EC project to the hippocampal DG, CA3, and CA2 (4-6). Pyramidal cells in layer III of the EC directly project to the hippocampal CA1(4, 7). A subpopulation of Wolfram syndrome 1 (Wfs1) / CalbindinD-28K (CalB)^+^ pyramidal cells in layer II of the EC project to the inhibitory neurons in the hippocampal CA1 area (4, 8, 9). On the other hand, in contrast to the superficial layers, the cell-type specific projection pattern for the deep layers of the EC are beginning to be studied. Layer V can be separated into two sublayers: Va and Vb (4, 9, 10). Vb neurons are known to function as a local projection to superficial layers in the EC (9, 11, 12), and Va neurons project to telencephalic structures (4, 9-11). Previous studies raised the possibility that some neurons in the deep layers of the EC also may project back to the HPC (13, 14). Utilizing molecular markers for deep layers and viral-based neural tracing, two recent studies revealed that Va neurons collaterally project to telencephalic structures as well as the hippocampal CA1 area (15). Furthermore, VI neurons project to the hippocampal CA1, CA2, CA3, and DG areas (16). However, it remains unknown whether Vb neurons in the EC also project to the HPC.

In this study, we investigated whether Vb neurons in the medial EC (MEC) project to the HPC. We injected retrograde tracers, Cholera Toxin Subunit B (CTB) 488 and CTB555, into the dorsal (AP:-2.00, ML:1.30, DV:-2.00) (**Fig. 1A**) and ventral DG (AP:-3.70, ML:2.90, DV:-3.50) (**Fig. 1B**) of five 6-10 weeks old C57BL6J male mice (JAX:000664), respectively (dDG: 100nl, vDG: 70nl, 0.5% wt/vol, Invitrogen). Six days after the injection, these mice were deeply anesthetized with a cocktail of ketamine (75mg/kg)/dexmedetomidine (1mg/kg) and then transcardially perfused with 4% paraformaldehyde (PFA) in PBS. Brains were extracted and post-fixed overnight in 4% PFA in PBS at 4 °C and then sectioned at a thickness of 60 μm using a vibratome (Leica). We found both CTB488^+^ and CTB555^+^ cells in layer II of MEC (MECII); CTB488^+^ cells were observed in the dorsal part of MECII while CTB555^+^ cells were found in the ventral part of MECII (**Fig. 1C-D, F**), as previously demonstrated (2, 3, 5), indicating that we successfully injected these tracers into the dorsal and ventral DG. We also found CTB488^+^ and CTB555^+^ cells in the deep layers of the MEC (**Fig. 1C-D, F-H**). There was a little overlap between CTB488^+^ and CTB555^+^ cells (**Fig. 1C-D, G-H**). To identify the cell-type of these cells, we examined them using immunohistochemistry for Ctip2, a marker for Vb neurons (9) (rat anti-Ctip2 antibody; ab18465, abcam, 1/300 dilution) and found that both CTB488^+^ and CTB555^+^ cells were colocalized with Ctip2 (**Fig. 1E-H**). Furthermore, we found that only the outer layer of Vb neurons was labeled with CTBs (**Fig. 1F-H**). We also found that CTB488^+^ cells were located in the dorsal part of MEC, while CTB555^+^ cells were more abundant than the CTB488^+^ cells and distributed through the dorso-ventral axis (**Fig. 1C-D, F**), indicating that neurons in the outer layer of Vb preferentially project to both dorsal and ventral DG (vDG), with a significant preference for the vDG. These CTB^+^ cells were not observed in Va (**Fig. 2A-L**). Since Va neurons project to the basolateral amygdala (BLA) and nucleus accumbens (NAc) (4, 8, 9), we could label Va neurons with CTB when we injected CTB into the NAc and BLA (**Fig. 1B-C, H-I**). We injected CTB488, CTB555, and CTB647 into the BLA (AP:-1.40, ML:3.40, DV:-5.00), vDG (AP:-3.70, ML:2.90, DV:-3.50), and NAc (AP:1.15, ML:0.70, DV:-4.70) of four 6-10 weeks old C57BL6J male mice, respectively (BLA 100nl, vDG: 70nl, NAc: 300nl, 0.5% wt/vol, Invitrogen). We found that vDG-projecting cells were never colocalized with BLA-or NAc-projecting cells, indicating that Va neurons do not project to the vDG.

**Figure 1.**
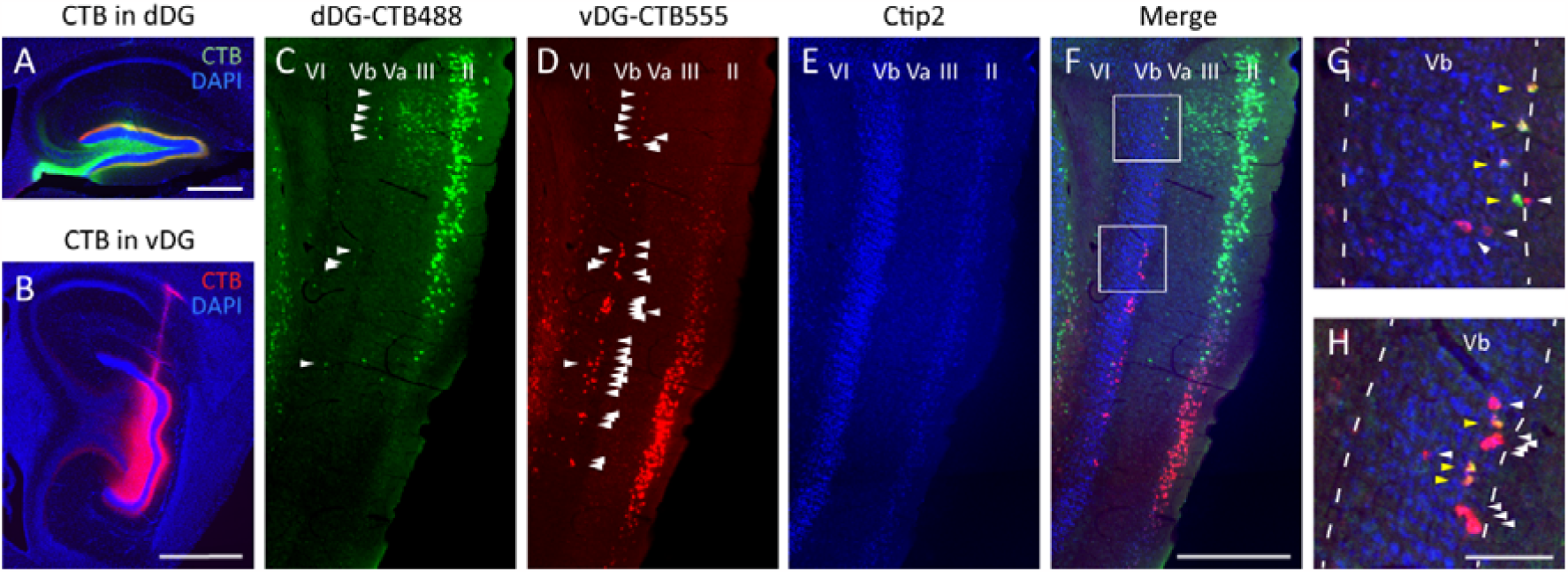
Outer layer of Vb neurons in medial entorhinal cortex project to hippocampal dentate gyrus in mice. **(A-B)** Injection of CTB488 and CTB555 into dorsal (A) and ventral DG (B), respectively. CTB488 (green). CTB555 (red). DAPI (blue). **(C-F)** Parasagittal sections of the MEC labeled with CTB488 (green), CTB555 (red) and immunostained with Ctip2 (blue). Arrowheads indicate CTB488^+^ cells (C) and CTB555^+^ cells (D) in Vb, individually. **(G-H)** Magnified images of F, top square (G) and bottom square (H), respectively. Yellow arrowheads indicate CTB488 and CTB555 double positive cells. White arrowheads indicate CTB555 single positive cells. Scale bar, 500μm in (A and F), 1mm in (B), and 100μm in (H).

**Figure 2.**
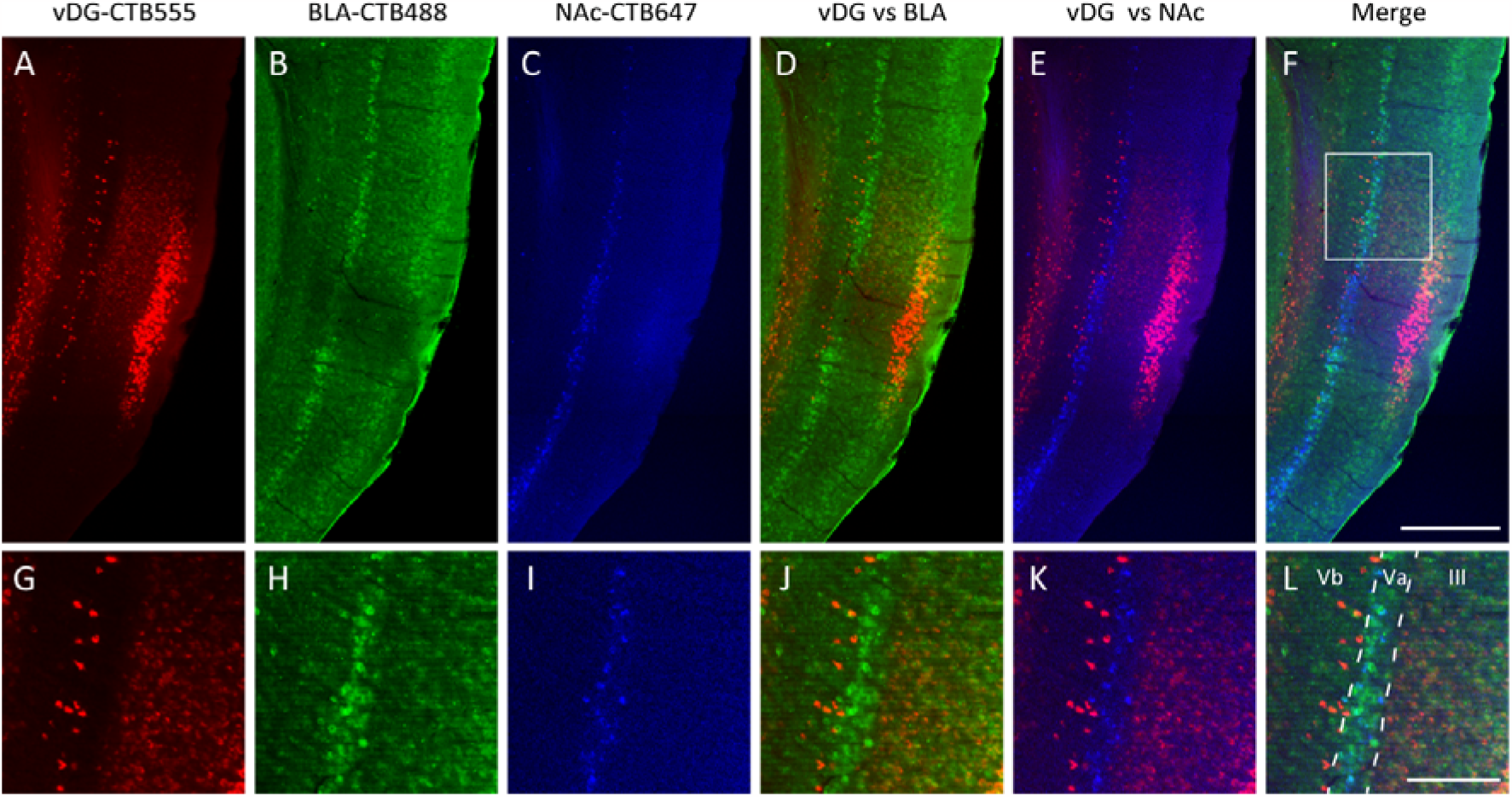
Difference of projection specific retrograde labeling in medial entorhinal cortex Va and Vb neurons. **(A-F)** Parasagittal sections of the MEC labeled with CTB488 (green), CTB555 (red), CTB647 (blue), which CTB488, CTB555, CTB647 were injected into BLA (B), vDG (A), and NAc (C), respectively. Scale bar, 500μm. **(G-L)** Magnified images of A-F, respectively. Scale bar, 200μm.

A previous study (9) showed that a subpopulation of Vb neurons projects to the anterior thalamic nuclei in mice. Therefore, to examine if DG-projecting Vb cells belong to the same population as anterior thalamus-projecting Vb cells, we injected CTB555 into the anterodorsal/anteroventral thalamic nucleus (AD/AV) (AP:-0.90, ML:0.90, DV:-3.15) (150nl, 0.5% wt/vol, Invitrogen) of eight 6-10 weeks old C57BL6J male and female mice (**Fig. 3A**). I examined the immunohistochemistry for Purkinje cell protein 4 (PCP4), a marker for ECIII and ECVb (10), (rabbit anti-PCP4, Sigma, HPA005792, 1/300) (**Fig. 3C**) and NeuN (chick anti-NeuN, abcam, ab134014, 1/1000) (**Fig. 3D**). In contrast to the DG-projecting Vb cells, which are located only in the outer layer of Vb (**Fig. 1F-H**), anterior thalamus-projecting Vb cells were distributed from the outer to the inner layer of Vb (**Fig. 3B, E-G**). Furthermore, double CTB injections into same mice (CTB488 in AD/AV and CTB555 in vDG, respectively, five 6-10 weeks old C57BL6J male and female mice) demonstrated that there is no overlap between vDG-projecting cells and AD/AV-projecting cells in Vb (**Fig. 3H**, examined total 171 vDG-projecting cells and 213 AD/AV-projecting cells from 5 mice, 0% overlap). These results suggests that DG-projecting Vb cells are different population from anterior thalamus-projecting Vb cells.

**Figure 3.**
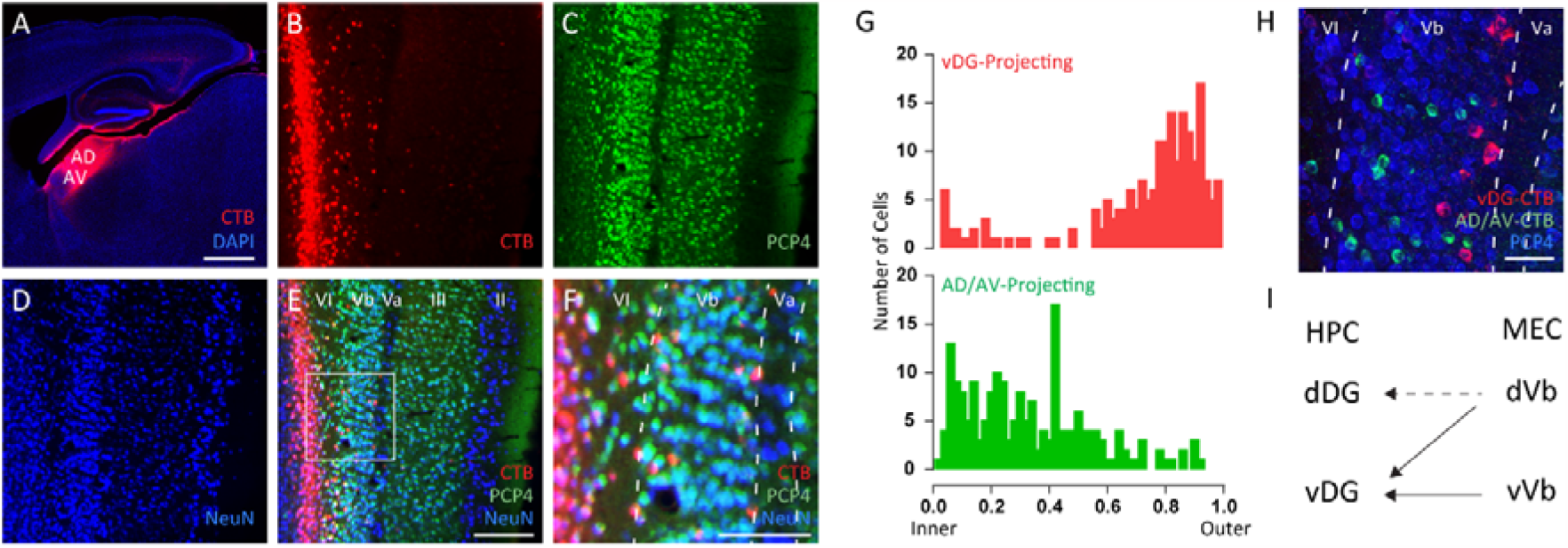
Distinct population of anterior thalamus-projecting and vDG-projecting Vb neurons in medial entorhinal cortex. **(A)** Injection of CTB555 into AD/AV. Scale bar, 1mm. **(B-E)** Parasagittal sections of the MEC labeled with CTB555 (red) and immuno-stained with PCP4 (green) and NeuN (blue). Scale bar, 200μm. **(F)** Magnified images of E. Scale bar, 100μm. **(G)** Distribution of vDG-projecting CTB^+^ cells (top, n = 7 mice) and AD/AV-projecting CTB^+^ cells (bottom, n = 6 mice) in MEC Vb which represented by 50 bins through the outer to the inner layer of Vb. **(H)** Parasagittal sections of the MEC Vb labeled with CTB488 (green) and CTB555 (red) and immune-stained with PCP4 (blue), which CTB488 and CTB555 were injected into AD/AV and vDG, respectively. Image was taken by confocal microscope. Scale bar, 50μm. **(I)** Summary for projection of MEC layer Vb neurons into hippocampus.

Vb neurons have been known to function as local projections to superficial layers in the EC to form the EC-HPC loop circuit (4, 9, 11, 12). However, in this study, we identified that the outer layer of MEC Vb neurons directly project to both the dorsal and ventral DG, with a significant preference for the ventral DG. DG-projecting Vb cells (**Fig. 3I**) and anterior thalamus-projecting Vb cells are distinct populations and they have differential distribution patterns in Vb (**Fig. 3G**). Previous studies showed that some layer V/VI cells in the EC project to the hippocampal DG (13, 14), but due to the lack of molecular markers for V/VI, the cell types remained unknown at that time. In this study, we revealed that outer layer of Vb neurons projects to hippocampal DG. Although it has long been considered that only superficial layers of EC neurons project to the HPC, accumulating evidence (15, 16) including this study indicates that the subpopulation of neurons from all layers in the EC differentially provide significant projections to the HPC. Further studies will be required to understand the neural processes and the computations underlying learning and memory, based on the updated anatomical maps in the EC-HPC networks.

## List of Abbreviations

AAV: Adeno-associated virus
AD: Anterodorsal thalamic nucleus
AV: Anteroventral thalamic nucleus
BLA: Basolateral amygdala
CalB: CalbindinD-28K
CTB: Cholera Toxin Subunit B
Ctip2: Chicken ovalbumin upstream promoter transcription factor (COUP-TF) interacting protein 2
DAPI-4′: 6-diamidino-2-phenylindole
dDG: dorsal Dentate gyrus
DG: Dentate gyrus
EC: Entorhinal cortex
HPC: Hippocampus
MEC: Medial entorhinal cortex
NAc: Nucleus accumbens
PCP4: Purkinje cell protein 4
vDG: ventral Dentate gyrus
Wfs1: Wolfram syndrome 1

## Declarations

### Ethics approval and consent to participate

All animal experiments were approved by the UT Southwestern Medical Center IACUC (Protocol# 2017-102301).

### Consent for publication

NA

### Availability of data and materials

Data for analysis will be made available by the corresponding author upon reasonable request.

### Competing interests

The authors declare that they have no competing interests.

### Funding

This work was supported by grants from Endowed Scholar Program (T.K), Human Frontier Science Program (T.K), Faculty Science and Technology Acquisition and Retention Program (T.K), and The Whitehall Foundation (T.K).

### Authors’ contributions

TK and SKO conceived the study. TK and SKO designed experiments. NY, JY, KR and SKO conducted experiments. JY and SKO analyzed data. NY, JY, TK and SKO interpreted data. TK and SKO wrote the manuscript.

## Acknowledgements

We thank all members of the Kitamura laboratory for their support.

## Notes

### Competing Interest Statement

The authors have declared no competing interest.

